# Enhanced population coding for rewarded choices in the medial frontal cortex of the mouse

**DOI:** 10.1101/429852

**Authors:** Michael J. Siniscalchi, Hongli Wang, Alex C. Kwan

**Affiliations:** Interdepartmental Neuroscience Program, Yale University School of Medicine, New Haven, Connecticut, 06511; Department of Psychiatry, Yale University School of Medicine, New Haven, Connecticut, 06511; Department of Neuroscience, Yale University School of Medicine, New Haven, Connecticut, 06511

**Keywords:** Choice history, ensemble activity, goal-directed actions, secondary motor cortex, two-photon calcium imaging

## Abstract

Instrumental behavior is characterized by the selection of actions based on the degree to which they lead to a desired outcome. However, we lack a detailed understanding of how rewarded actions are reinforced and preferentially implemented. In rodents, the medial frontal cortex is hypothesized to play an important role in this process, based in part on its capacity to encode chosen actions and their outcomes. We therefore asked how neural representations of choice and outcome might interact to facilitate instrumental behavior. To investigate this question, we imaged neural ensemble activity in layer 2/3 of the secondary motor region (M2) while mice engaged in a two-choice auditory discrimination task with probabilistic outcomes. Correct choices could result in one of three reward amounts (single-, double-, or omitted-reward), which allowed us to measure neural and behavioral effects of reward magnitude, as well as its categorical presence or absence. Single-unit and population decoding analyses revealed a consistent influence of outcome on choice signals in M2. Specifically, rewarded choices were more robustly encoded relative to unrewarded choices, with little dependence on the exact magnitude of reinforcement. Our results provide insight into the integration of past choices and outcomes in the rodent brain during instrumental behavior.

## Introduction

Associations between past choices and their outcomes allow for efficient selection of actions likely to meet one’s present goals. The mechanisms through which such associations are implemented remain unclear. However, at the behavioral level it is well known that rewarded choices tend to be repeated at the expense of those that have yielded meager or aversive results. To gain insight into the associative mechanisms underlying goal-directed action selection, the present study focuses on the question of how populations of simultaneously recorded neurons in the cerebral cortex represent and integrate information related to choices and their outcomes.

How does the brain selectively reinforce rewarded actions in order to bias their future implementation? Physiological studies in primates and rodents suggest that the frontal lobe plays an important role in these functions. For example, the primate prefrontal cortex is known to contain neurons that encode chosen actions and outcomes (Barraclough DJ et al. 2004; Genovesio A et al. 2006; Seo H et al. 2007; Histed MH et al. 2009), suggesting a plausible neural substrate for their association during goal-directed behavior. Moreover, single-unit recordings have revealed that prior reward enhances the discriminability of spiking activity related to past (Donahue CH et al. 2013) and upcoming choices (Histed MH *et al*. 2009).

Similarly, recordings from the medial frontal cortex (MFC) of rodents have revealed neural signatures of prior choices and outcomes (Sul JH et al. 2010; Sul JH et al. 2011; Hyman JM et al. 2017). In particular, these studies have demonstrated neural representations of choice and outcome history in the most dorsal anatomical sub-region of MFC, which is referred to as secondary motor cortex (M2) in mice, and medial agranular or medial precentral cortex in rats (Sesack SR et al. 1989; Barthas F and AC Kwan 2017). Murine M2 has also been implicated in the flexible acquisition and initiation of voluntary actions (Ostlund SB et al. 2009; Gremel CM and RM Costa 2013; Murakami M et al. 2014; Siniscalchi MJ et al. 2016; Barthas F and AC Kwan 2017; Makino H et al. 2017). Based on its putative role in instrumental behavior, M2 may serve as an important interface for the mixing of choice- and reward-related signals in the rodent brain. However, the details of how reinforcement might interact with choice-related neural representations remains unclear. One intriguing hypothesis is that a choice’s outcome could affect the strength or persistence of its neural signature in M2.

To explore this possibility, we trained mice on a two-choice auditory discrimination task, and then introduced a probabilistic reinforcement schedule during testing. Simultaneous two-photon calcium imaging enabled the characterization of task-related neural ensemble activity in M2. Three randomly interleaved outcomes (single-, double-, and omitted- rewards) delivered following correct responses allowed us to measure the impact of reward on choice coding, as well as to distinguish effects of changes in reward magnitude from those of its absolute presence or absence. We found that rewarding outcomes boosted the fidelity of choice signals encoded in the population activity patterns—an effect that persisted into the subsequent trial. Importantly, the reward-dependent enhancement of choice-related signals depended less on differences in reward size than on the categorical presence or absence of rewards. These results suggest one plausible cortical mechanism for the reinforcement of rewarded actions—namely, that rewarding outcomes lead to a more robust population level read-out of recent choice history.

## Materials and Methods

### Animals

All procedures were performed in accordance with the regulations of the Institutional Animal Care and Use Committee at Yale University. Animals were housed on a 12-h/12-h light-dark cycle (lights off at 19:00) in groups of three-to-five per cage. Ten adult (postnatal day 111 – 279) male mice with a C57BL/6J (#000664, Jackson Laboratory) genetic background were used. Although two subjects from these experiments were also used in an earlier study (Siniscalchi MJ *et al*. 2016), the data and analyses in this paper have not been reported elsewhere.

### Surgery

For optical imaging of neural activity, we implanted a chronic glass window over the medial secondary motor cortex (M2). The surgical procedures were nearly identical to those used in our earlier studies (Phoumthipphavong V et al. 2016). All subjects were treated pre-operatively with carprofen (5 mg/kg, s.c.; #024751, Butler Animal Health) and dexamethasone (3 mg/kg, s.c.; Dexaject SP, #002459, Henry Schein Animal Health). Anesthesia was then induced with 2% isoflurane in oxygen, and the animal was placed on a water-circulating heating pad (TP-700, Gaymar Stryker). Following induction, isoflurane concentration was lowered to 1.5%, and the head was secured in a stereotaxic frame with ear bars (David Kopf Instruments). The scalp was shaved with electric trimmers, and cleaned with a povidone-iodine surgical scrub (Betadine, Perdue Products L.P.). A narrow portion of scalp was removed along the midline from the interaural line to a line visualized just posterior to the eyes. The incision was retracted to expose the dorsal aspect of the skull, which was then scrubbed briefly with 3% hydrogen peroxide to aid in removal of the periosteum, and washed generously with artificial cerebrospinal fluid (ACSF; in mM: 5 KCl, 5 HEPES, 135 NaCl, 1 MgCl_2_, and 1.8 CaCl_2_; pH 7.3). A 3-mm-diameter circular craniotomy was made over the right hemisphere using a high-speed rotary drill (K.1070, Foredom), centered on a medial target within M2 (AP +1.5 mm, ML –0.5 mm relative to Bregma). The dura was left intact, and was irrigated frequently with ACSF over the remainder of the procedure. An adeno-associated virus encoding GCaMP6s (AAV1-Syn-GCaMP6s-WPRE-SV40, Penn Vector Core) was infused at four AP-ML coordinates through a glass micropipette attached to a microinjection unit (Nanoject II, Drummond). The injection sites formed a 200-um-wide square centered on the target location. Each site was injected with 46 nL of virus over 3 min, at a depth of 0.4 mm. To minimize backflow of the injected solution, the micropipette was left in place for 5 min after each infusion. A small amount of warmed agarose solution (Type III-A, High EEO agarose; 1.2% in ACSF; #A9793, Sigma Aldrich) was then applied along the perimeter of the craniotomy to fill the space between the cranial window and the surrounding tissue after implantation. The cranial window consisted of two concentric circular glass parts, glued together with UV-activated optical adhesive (NOA 61, Norland Products, Inc.): a 3-mm-diameter, #1 thickness prefabricated glass coverslip (#64–0720-CS-3R, Warner Instruments), and a 2-mm-diameter plug cut from a #1 or #2 thickness glass coverslip. The window was carefully placed on the brain surface with the glass plug facing down, and then secured to the skull at the edge of the craniotomy using cyanoacrylate glue. Gentle downward pressure was applied to stabilize the implant during this procedure, using a wooden probe attached to the stereotaxic frame. A custom-made stainless-steel head plate (eMachineShop) was then bonded to the skull with dental cement (C&B Metabond, Parkell Inc.), with care taken to cover any remaining exposed skull. Post-operative care was provided immediately, and for three consecutive days following surgery. This consisted of analgesia (carprofen, 5 mg/kg, s.c.) and fluid support (Preservative-free 0.9% NaCl, 0.5 mL, s.c., up to twice daily). All animals were given a one-week post-operative recovery period prior to the onset of behavioral training.

### Behavioral setup

Subjects were placed in a modified acrylic tube (8486K33, McMaster-Carr), and held head-fixed during the behavioral task by fastening the head plate implant to a custom-made stainless-steel bracket (eMachineShop). This setup limits gross body movements, but allows for small postural adjustments and movement of the hind limbs. Two lick spouts, mounted on a 3D-printed plastic part, were placed on either side of the mouth to allow lateralized responses and corresponding delivery of water rewards. This basic two-choice setup was modeled after an earlier study (Guo ZV et al. 2014). Lick spouts were fabricated from 20-guage hypodermic needles, which were cut and carefully filed with an abrasive wheel, and then soldered to a wire lead and connected to a battery-operated lick detection circuit (Slotnick B 2009). Output signals from the detector were digitized with a USB data acquisition device (USB-201, Measurement Computing) plugged into a desktop computer. Water rewards were delivered through each spout by gravity-feed, and actuated with a solenoid valve (EV-2–24, Clippard) controlled by TTL pulses from a second USB-201. Pulse duration for single-reward (2 μL) was calibrated for each valve by sending 100 pulses and then weighing the ejected volume (a 15 – 20 ms TTL pulse was typically required). On double-reward trials, 4 μL were delivered by using either a calibrated longer-duration pulse, or two single-rewards separated by 100 ms. Auditory stimuli were played through a pair of computer speakers (S120, Logitech) placed in front of the animal. The task structure was automated using custom scripts written for Presentation (Neurobehavioral Systems, Inc.). During training, the behavioral apparatus was enclosed in the cabinet of an audiovisual cart (4731T74, McMaster-Carr) soundproofed with acoustic foam (5692T49, McMaster-Carr). For imaging, the setup was replicated within the enclosure of a two-photon microscope.

### Two-choice auditory discrimination task with probabilistic outcomes

Mice were trained to perform a two-choice auditory discrimination task. To motivate participation, subjects were water restricted, as follows. Six days per week, water was provided only in a single daily training session, as a reward for correct choices. On the remaining day, a water bottle was placed in the home cage for fifteen minutes. Three phases of training were used to shape the behavior, identical to those used in our prior study (Siniscalchi MJ *et al*. 2016). During phase one, mice were habituated to head fixation and trained to lick the left or right spout for water: each lick to either spout triggered the release of 4 μL of water, with a minimum interval of one second between rewards. After attaining > 100 rewards in a single session of phase one (1–2 days), subjects were advanced to phase two, in which they were required to lick for a similar number of rewards at each spout. In this case, water was only available from one ‘target’ spout at a time, with the target moving to the opposite side each time the mouse earned three consecutive rewards from a given target. Additionally, sound stimuli were introduced in association with rewarded licks (‘hits’) on a given spout. The stimuli were trains of four 500-ms-long logarithmic chirps, with starting and ending frequencies of either 5 and 15 kHz (‘upsweep’), or 15 and 5 kHz (‘downsweep’), respectively. Upsweeps were played immediately following a left hit, and downsweeps were played immediately following a right hit. Similar to phase one, a minimum interval of one second was imposed between rewards. After attaining > 100 hits within a single session of phase two (1–2 days), subjects were advanced to phase three, in which they were trained to perform a two-choice sound discrimination task. Unlike the earlier two phases, in which the operant behavior was self-paced, phase three was structured into trials with a defined response period. Each trial began with the presentation of a 2-s-long sound cue (upsweeps or downsweeps, randomly drawn), which indicated the target spout for that trial. Upsweeps indicated ‘left’ and downsweeps indicated ‘right’. To earn a water reward, the subject was required to lick the target spout within the final 1.5 s of the cue (the ‘response window’). The first lick to either spout within the response window terminated the sound cue and triggered an immediate outcome: 2 μL of water from the target spout for a hit, or playback of a 2-s-long white noise sound for an ‘error’, in which the wrong spout was chosen. ‘Misses’ were defined as trials in which the mouse failed to lick within the response window. In any case, the next trial would begin 7 s after cue offset. Thus, trial durations ranged from 7.5 to 9 s, depending on response time. Each session was terminated automatically after twenty consecutive misses. For the current study, subjects were trained on the two-choice sound discrimination task to high level of proficiency (>90% hit rate), and were then tested on a task variant with probabilistic outcomes. This variant was identical to phase three of training, except that correct responses could result in one of three outcomes: 2 μL of water (‘single-reward’) with 80% probability, no reinforcement (‘omitted-reward’) with 10% probability, or 4 μL of water (‘double-reward’) with 10% probability. In a subset of sessions (five out of sixteen, identified in **Table 1**), the white noise sound used in error trials was also played at the time of an omitted reward. We analyzed the behavioral and neural data for those sessions separately, but did not detect any obvious differences; therefore, we have presented the pooled results.

**Table 1:** Mice used in this study.

| Experiment | Exp. number | Subject | Cells | Trials | Hit rate (%) | White noise in omitted-reward trial |
| --- | --- | --- | --- | --- | --- | --- |
| M2 imaging | 1 | L4 | 35 | 198 | 99.0 | ✓ |
| M2 imaging | 2 | L2 | 47 | 228 | 80.7 | ✓ |
| M2 imaging | 3 | M6 | 62 | 321 | 93.2 | ✓ |
| M2 imaging | 4 | M6 | 42 | 233 | 97.9 | ✓ |
| M2 imaging | 5 | M6 | 55 | 212 | 95.3 | ✓ |
| M2 imaging | 6 | M12 | 51 | 336 | 94.9 |  |
| M2 imaging | 7 | M20 | 46 | 182 | 91.8 |  |
| M2 imaging | 8 | M12 | 40 | 300 | 95.7 |  |
| M2 imaging | 9 | M13 | 43 | 313 | 93.0 |  |
| M2 imaging | 10 | M12 | 44 | 285 | 98.3 |  |
| M2 imaging | 11 | M14 | 38 | 210 | 99.1 |  |
| M2 imaging | 12 | M16 | 59 | 202 | 88.6 |  |
| M2 imaging | 13 | M16 | 77 | 269 | 93.7 |  |
| M2 imaging | 14 | M17 | 49 | 267 | 91.8 |  |
| M2 imaging | 15 | M17 | 33 | 222 | 92.8 |  |
| M2 imaging | 16 | M22 | 50 | 262 | 94.3 |  |

### Two-photon calcium imaging

The two-photon microscope (Movable Objective Microscope, Sutter Instrument) was controlled using ScanImage software (Pologruto TA et al. 2003). The excitation source was a Ti:Sapphire ultrafast femtosecond laser (Chameleon Ultra II, Coherent). Beam intensity was modulated using a Pockels cell (350–80-LA-02, Conoptics) and blanked with an optical shutter (LS6ZM2, Uniblitz / Vincent Associates). The beam was focused through a high-numerical aperture objective (XLUMPLFLN, 20X/0.95 NA, Olympus). Excitation wavelength was set at 920 nm, and the emission was collected behind a bandpass filter from 475 to 550 nm using a GaAsP photomultiplier tube (H7422P-40MOD, Hamamatsu). The time-averaged excitation intensity after the microscope objective was ∼100 mW. Time-lapse image frames were acquired at 256 × 256 pixels, with a frame rate of 3.62 Hz using bidirectional raster scanning. To synchronize the behavioral and imaging data, a TTL pulse was sent by Presentation one second prior to the start of each trial. This TTL signal was assigned as an external trigger in ScanImage to initiate a new image file. Timestamps of the TTL pulses along with timestamps of other behavioral events were written to a text file by Presentation, allowing the image files to be aligned with behavioral events.

### Analysis of behavioral data

Timestamps of the behavioral events, including cue onsets, licks, and reinforcement onsets, were logged to a text file by Presentation. All further processing and analysis was done using custom scripts written in MATLAB (MathWorks, Inc.). The number of trials performed was defined as the number of trials in which either spout was licked within the response window. Correct rate was defined as the number of correct trials divided by the number of trials performed. Miss rate was defined as the number of misses divided by the total number of trials. The sensitivity index, or d-prime, was calculated as the difference between the inverse of the standard normal cumulative distribution for the correct rate on upsweep trials and the inverse of the standard normal cumulative distribution for the incorrect rate on downsweep trials.

### Analysis of imaging data

Raw time-lapse image stacks corresponding to each trial were saved as multipage TIFF files by ScanImage. As a first processing step, the raw stacks were merged to a single TIFF. Timestamps for the first frame of each trial, as well as the external trigger, were extracted for alignment with behavior. The merged TIFF file was then processed for *x*-*y* motion correction using either the TurboReg (Thevenaz P et al. 1998) or moco (Dubbs A et al. 2016) plug-in for ImageJ (Schneider CA et al. 2012). Regions of interest (ROIs) were manually selected around cell bodies appearing in the maximal or average projection image, using a custom graphical user interface programmed in MATLAB. Pixel intensity values within each ROI were summed for each frame *t* to obtain *F*(*t*). The baseline fluorescence, *F_o_*(*t*), was estimated as the 10^th^ percentile of *F*(*t*) within a 10 min moving window centered on *t*. Δ*F*/*F*(*t*) was then calculated as the fractional change in *F*(*t*) relative to *F_o_*(*t*).

### Event-aligned activity and choice selectivity

To obtain trial-averaged activity traces associated with a specific behavioral event (e.g., cue onset), we first aligned Δ*F*/*F*(*t*) traces based on their timing relative to each instance of the event, and then took the mean across traces. To estimate confidence intervals, we performed a bootstrapping procedure, as follows. For *N* instances of a particular event, *N* traces were resampled randomly with replacement and then averaged over 1000 iterations, in order to approximate the sampling distribution for the mean. Upper and lower bounds of the confidence interval were then estimated as percentiles of this distribution. Choice selectivity was calculated as the difference between cue-aligned trial-averaged traces from trials in which the ipsilateral versus contralateral spout was chosen, divided by the sum of the two traces. Therefore, choice selectivity was a function of time that could take values from −1 to 1, with negative values indicating an ipsilateral preference and positive values indicating a contralateral preference. The mean choice selectivity from 2 to 4 s after cue onset was used as a scalar estimate in comparisons between double- and omitted-reward trials.

### Multiple linear regression

To characterize the relationship between task variables and the activity of individual neurons, we fit the following linear equation: 
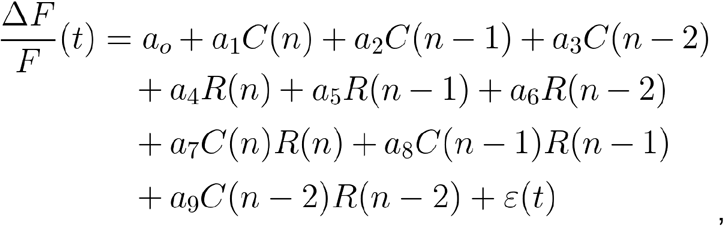

where Δ*F*/*F*(*t*) is the fractional change in fluorescence at time *t* relative to baseline; *C*(*n*), *C*(*n*-1), and *C*(*n*-2) are the choices made on the current trial, the prior trial, and the trial before last; *R*(*n*), *R*(*n*-1), and *R*(*n*-2) are the outcomes for the current trial, the prior trial, and the trial before last; *a_1…_a_9_* are the regression coefficients, and *ε*(*t*) is the error term. Choices were dummy-coded as −1 for left licks and 1 for right licks. Outcomes were dummy-coded as 0 for single-reward trials, −1 for omitted-reward trials, and 1 for double-reward trials. Error trials were ignored in the analysis (set to NaN in MATLAB). Regression coefficients and their p-values were estimated for each 500 ms time bin *t*, within an interval from −2 s to 6.5 s relative to cue onset. To summarize the pooled results from all experiments, the proportion of neurons with a p-value less than 0.01 was plotted over time for each predictor. The binomial test was used to determine whether this proportion was significantly different from chance level.

### Decoding: linear discriminant analysis

To determine how reliably the subject’s choice was encoded in the neural ensemble activity, we constructed and tested classifiers based on linear discriminant analysis. All classifiers were constructed using the ‘classify’ function in MATLAB, with the ‘type’ parameter set to ‘linear’ (the default); this classification method fits a multivariate normal density function to each group, using a pooled estimate of covariance. Choices, *C*(*n*), were dummy-coded as −1 for left licks and 1 for right licks. For each 500 ms time bin *t* within the interval from −2 s to 6.5 s relative to cue onset, the Δ*F*/*F*(*t*) values of all neurons were incorporated into a trial-indexed set of population activity vectors, with each vector representing the Δ*F*/*F*(*t*) for all neurons in the corresponding trial. A Monte Carlo cross-validation procedure was then applied across trials, as follows. First, activity vectors from a randomly drawn 80% of single-reward trials were used as the training set to construct a classifier. The classifier was then tested for accuracy using the activity vectors from the remaining 20% of single-reward trials. Additionally, we tested the accuracy of the classifier at decoding choice in other trial types, i.e. double-reward, omitted-reward, and error trials. To estimate chance-level performance for each classifier, an identical classifier was constructed and tested, with the exception that the *C*(*n*) values within the training set were first shuffled randomly. To characterize the potential advantage of simultaneous recording, we compared the accuracy of classifiers trained on actual imaging data versus ‘pseudo-ensemble’ data in which simultaneity had been destroyed. To build a pseudo-ensemble, the Δ*F*/*F*(*t*) values for each cell were randomly shuffled across all training trials with the same *C*(*n*) value. Therefore, each pseudo-ensemble activity vector comprised Δ*F*/*F*(*t*) values drawn from many different trials, while preserving cell identity as well as choice and outcome specificities. Classifiers trained on pseudo-ensemble data versus real data were then compared for accuracy using the remaining (unshuffled) test trials. For all of the analyses described in this section, average decoding accuracy was estimated as the mean across 30 iterations. This iterative cross-validation procedure was repeated for each time bin *t*.

### Decoding: random forests

As a second approach to decoding choices from the population activity, we constructed and tested random forest classifiers (Breiman L 2001). Neural and behavioral data were treated in the same manner as for the linear discriminant analysis described above: choices, *C*(*n*), were dummy-coded as −1 and 1; time ranged in 500 ms increments from −2 to 6.5 s relative to cue onset; and single-reward trials were split into training and testing sets for 30 iterations of Monte Carlo cross-validation. For each iteration, a random forest classifier was trained using population activity vectors from a random sample of 80% of trials, and then tested on the remaining 20%. The random forest algorithm is a bootstrap aggregation (‘bagging’) approach consisting of many decision trees. Each decision tree takes as input a set of features (e.g., the activity of cells in the neural ensemble) and arrives at a predicted binary response (e.g., left or right choice). It does so by comparing a subset of the features to a series of corresponding threshold values (‘splits’) learned from the training set through a process of greedy recursive partitioning. The overall predicted response of the random forest is the majority vote of the predicted responses across all trees. To construct each decision tree, *M* population activity vectors were drawn randomly with replacement from the *M* trials making up the training set. Each split in the tree comprised a threshold on the Δ*F*/*F*(*t*) value of one cell selected as the strongest predictor out of a randomly drawn subset of cells. A new subset was drawn randomly without replacement to determine each split. The number of cells in each subset was equal to the square root of *N*, where *N* is the total number of cells in the imaged ensemble. To choose the number of trees, we tested a range of values and found that classifier performance saturated beyond ∼50 trees; therefore, the number of trees was set to 100. The procedure was implemented by calling the ‘fitensemble’ function in MATLAB with the ‘Method’ parameter set to ‘Bag’, the ‘Type’ parameter set to ‘Classification’, and the ‘Learners’ parameter set to ‘Tree’. Additionally, in the same manner as we did for the linear discriminant analysis, we tested classifiers constructed using single-reward trials on other trial types, determined chance-level accuracy by training classifiers on shuffled *C*(*n*)’s, and characterized the advantage of simultaneous recording by comparing with random forest classifiers constructed using pseudo-ensemble data. For all decoding analyses described in this section, average decoding accuracy was estimated as the mean accuracy across all iterations. This entire procedure was repeated for each time bin *t*.

### Decoding: varying the ensemble size

To determine how the number of neurons in an ensemble influences decoding accuracy, we constructed and tested classifiers on neural data drawn from subsets of the imaged populations. Because the smallest ensemble imaged in our experiments had just over 30 neurons, we limited this analysis to ensemble sizes from 1 to 30 cells. We also limited the analysis to the time interval from 2 to 4 s from cue onset, the period in which our decoding analyses showed the highest decoding accuracy. For each ensemble size, a subset of the imaged cells was drawn randomly without replacement to produce an ensemble of predetermined size, and the process was repeated for 30 draws. For each draw, a classifier was constructed using a random subsample of 80% of trials, and then tested on the remaining trials. Decoding accuracy was estimated as the mean across draws. This iterative procedure was repeated for each ensemble size, and for the two types of classifiers (linear discriminant and random forest).

### Experimental Design and Statistical Analysis

The structure of the task was fully automated using custom scripts written for Presentation, which randomized the sound cues and outcomes presented in each trial. No further blinding was used. Sample sizes are noted in the results and figure legends. No statistical analysis was employed to determine sample sizes; however, they were similar to those used elsewhere in the field. All behavioral and neural ensemble analyses were performed using a sample size of *N* = 16 imaging sessions from ten animals (**Table 1**). For analyses of single unit activity (**Figs. 4–5**), the sample size was *N =* 771 cells; a subsample of *n* = 226 choice-selective cells was considered in Fig. 5 (see Results). A total of seven sessions were excluded from the study prior to neural activity analysis for the following reasons: poor behavioral performance (overall correct rate < 80%, one session); too few trials completed, and therefore fewer than five trials for at least one outcome type (three sessions); or residual movement artifacts after image motion correction (three sessions). All statistical analyses were performed in MATLAB (MathWorks, Inc.). A paired design was used for comparisons across outcome conditions, and the likelihood *p* of a false positive was estimated with a Wilcoxon signed-rank test. *p* < 0.05 was taken to indicate a significant difference. No corrections were made for multiple comparisons, but *p*-values are noted explicitly in the Results. For the multiple linear regression analysis (**Fig. 4**), the significance threshold for each predictor was set at *α* = 0.01. Significant proportions were determined using a binomial test, with *α* = 0.01. For neural ensemble analyses, chance-level accuracy of decoding choices from the neural activity was determined by testing classifiers constructed using shuffled choices.

## Results

### Two-choice discrimination task with probabilistic outcomes

Water-restricted mice were trained to perform a two-choice auditory discrimination task under head restraint (**Fig. 1A**). Subjects were required to choose between two lick spouts placed on either side of the mouth, only one of which (the ‘target’) would be rewarded on a given trial. One of two sound cues was presented at the start of each trial, indicating the target spout. The sound cues were trains of four 500-ms-long logarithmic chirps, with starting and ending frequencies of either 5 and 15 kHz (‘upsweep’), or 15 and 5 kHz (‘downsweep’), respectively. Upsweeps indicated ‘left’ and downsweeps indicated ‘right’. Licking either spout triggered an immediate outcome: delivery of a water reward if the target spout was chosen (‘correct’), or playback of a 2-s-long white noise sound if the wrong spout was chosen (‘error’). To study the influence of reward on behavior and neural activity, correct responses were reinforced probabilistically with one of three water amounts: 2 μL (‘single-reward’) with 80% probability, 4 μL (‘double-reward’) with 10% probability, or 0 μL (‘omitted-reward’) with 10% probability. In a subset of sessions, the white noise sound used in error trials was also played at the time of an omitted reward (**Table 1**). Each session was terminated automatically after twenty trials without a response (‘misses’).

**Figure 1.**
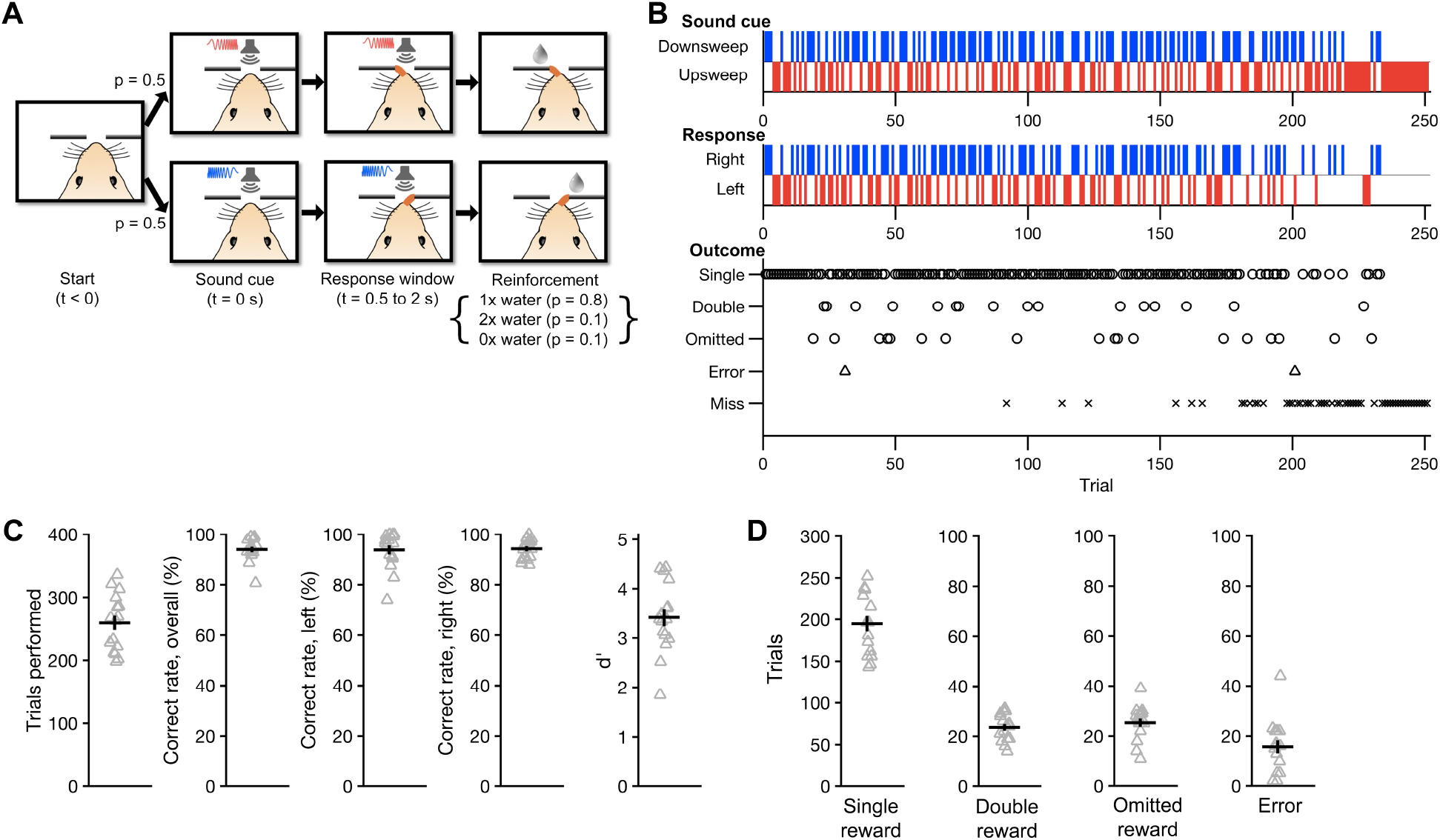
Two-choice auditory discrimination task with probabilistic outcomes. On each trial, mice were required to lick the target spout (left or right) indicated by a sound cue (upsweep or downsweep, respectively). Correct responses were rewarded probabilistically with one of three water amounts. ***A***, Flow diagram of the trial structure on correct trials. Each trial began with one of the two sound cues. The first lick to the target spout within 0.5–2 s following cue onset (Response window) immediately triggered one of three outcomes (Reinforcement): single-reward (1x), double reward (2x), or omitted-reward (0x), with probabilities of 80%, 10%, and 10%, respectively. The next trial would begin 7 s after cue offset. ***B***, Behavioral performance from an example session (Experiment 1 in Table 1). Occurrences of each sound cue (top), choice (middle), and outcome (bottom) are displayed in raster form according to trial number. Errors occurred when the non-target spout was chosen for the first lick within the response window. Misses were defined by the failure to respond within the response window. ***C***, Summary of behavioral performance. Gray triangles, individual sessions. Black crosshairs, mean ± *SEM. **D***, Number of occurrences of each outcome per session. For all figures, *N* = 16 sessions from 10 mice unless otherwise noted.

The use of probabilistic outcomes allowed us to systematically investigate neural and behavioral effects of reward during sensorimotor decision making. In particular, the inclusion of omitted-reward trials permitted the effects of reward absence to be explicitly characterized during correct trials, thus eliminating differences in decision accuracy as a potential confound. Additionally, the influence of reward size (single- vs. double-reward) could be compared to that of its categorical presence or absence (single- or double- vs. omitted-reward).

We obtained sixteen imaging sessions from ten mice while they performed this task (range: 1 – 3 sessions per mouse; **Table 1**). **Figure 1B** shows the behavioral performance from one such session. Subjects made 259 ± 11 choices per session, with an overall correct rate of 94 ± 1% (mean ± *SEM*; **Fig. 1C**), and a sensitivity index (*d’*) of 3.4 ± 0.2. Within each session, they encountered an average of 195 ± 9 single-reward, 24 ± 1 double-reward, and 25 ± 2 omitted-reward trials, and made 16 ± 3 errors. Correct rates were similar for upsweep (94 ± 2%) and downsweep (94 ± 1%) trials. Thus, subjects maintained a high level of accuracy following the introduction of variable outcomes.

### Outcome-dependent behavioral adjustments on multiple timescales

Subjects were well-trained in auditory discrimination before probabilistic outcomes were introduced, and the optimal stimulus-response relationships remained the same. Therefore, behavioral adjustments based on the new distribution of outcomes would not increase the overall amount of reward obtained. This raises an important question: were subjects aware of the different outcomes? Two lines of evidence indicate that they were. Firstly, the number of licks to the target spout during the period following reward delivery increased in a graded manner with the volume of water reward given (**Fig. 2A**). On average, 13.9 ± 0.8 licks were registered at the target spout in single-reward trials (mean ± *SEM, N* = 16 sessions). Relative to single-reward trials, the mean number of licks per trial rose by 21 ± 4% in double-reward trials (vs. single-reward trials: *p* = 0.001, Wilcoxon signed-rank test, *N* = 16 sessions), and decreased by 48 ± 3% in omitted-reward trials (vs. single-reward trials: *p* = 4 × 10^−4^, Wilcoxon signed-rank test, *N* = 16 sessions). Because the different outcome types were interleaved randomly across correct trials, these results indicate that the mice could detect changes in reward volume, and adjusted their consummatory licking accordingly.

**Figure 2.**
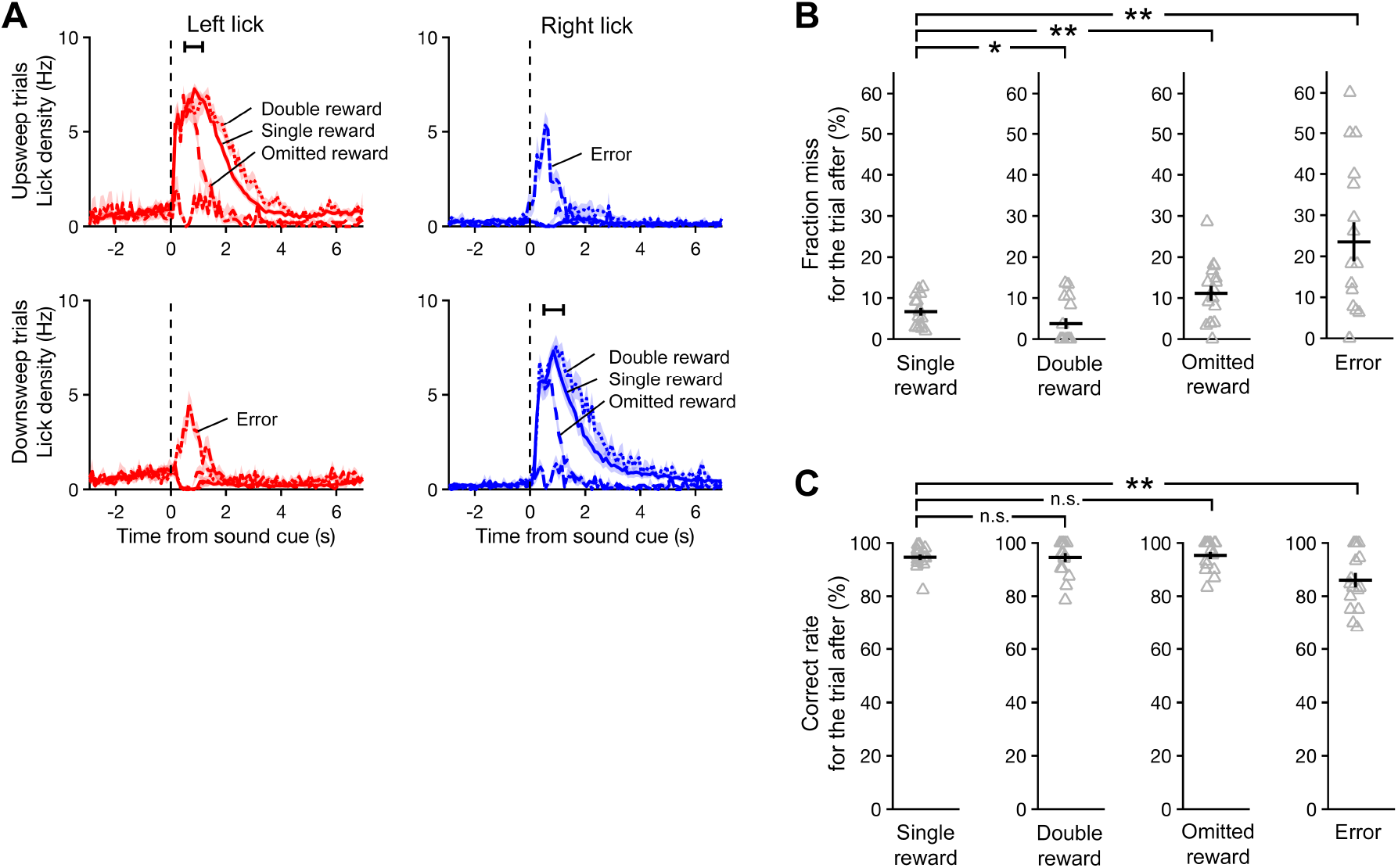
Subjects adjusted their behavior based on trial outcome. ***A***, Mean lick density across sessions as a function of time for the left and right spouts, averaged separately from trials in which the sound cue was an upsweep (top row) or downsweep (bottom row), and outcome was single- (solid), double- (dotted), or omitted-reward (dashed). Black error bar, 95% confidence interval for time of outcome. ***B***, Fraction of trials missed immediately following each outcome. Gray triangles, individual sessions. Black crosshairs, mean ± *SEM*. Wilcoxon signed-rank test: \**p* < 0.05, \*\**p* < 0.005, n.s., not significant. ***C***, Fraction of trials with a correct response immediately following each outcome.

Secondly, behavioral performance varied as a function of the prior trial’s outcome. The most notable effect was on the number of misses (**Fig. 2B**). The likelihood of a miss was 7 ± 1% for trials following a single reward (mean ± *SEM, N* = 16 sessions). It was significantly lower at 4 ± 1% for trials following a double reward (*p* = 0.04, Wilcoxon signed-rank test, *N* = 16 sessions), but significantly higher at 11 ± 2% if reward was omitted in the previous trial (vs. single-reward trials: *p* = 0.005, Wilcoxon signed-rank test, *N* = 16 sessions). However, when subjects did respond, the correct rate was consistently high at 95 ± 1%, 95 ± 2%, and 95 ± 1% for trials following a single, double, and omitted reward, respectively (**Fig. 2C**; mean ± *SEM, N* = 16 sessions). Taken together, the observed adjustments in licking and the effect on miss rate in the subsequent trial provide clear evidence that the mice monitored trial outcomes. Notably, the magnitude of reinforcement affected subjects’ willingness to respond, but did not influence the accuracy of decisions.

### Persistent decline in performance associated with errors

Errors were uncommon in these experiments. However, inference to the causes of residual inaccuracy could help to illuminate internal processes underlying decision-making. Our task design permitted us to examine and rule out two potential sources of error. Firstly, errors could have resulted mainly from stochastic fluctuations in perceptual performance. In this case, the likelihood of a correct response should not depend significantly on recent performance history. Thus, similar levels of performance would be expected in trials following errors and correct responses. However, relative to correct responses, errors were associated with deficits in both decision accuracy and willingness to respond in the subsequent trial. Correct rate dropped to 86 ± 3% in trials following an error (**Fig. 2C**; versus single-reward trials: *p* = 0.01; versus omitted-reward trials: *p* = 0.01, Wilcoxon signed-rank test, *N* = 16 sessions), and miss rate increased to 23 ± 5% (**Fig. 2B**; versus single-reward trials: *p* = 0.004; versus omitted-reward trials: *p* = 0.04, Wilcoxon signed-rank test, *N* = 16 sessions). We also asked whether the likelihood of an error could be explained by outcome-dependent processes, such as exploration following the absence of an expected reward. This hypothesis is not supported by the observation that error rates following correct trials were similar regardless of the prior trial’s outcome (**Fig. 2C)**. Collectively, our results suggest that the drop in task performance associated with errors tends to persist and cannot be solely accounted for by the trial outcome.

### Single-unit activity related to choices and outcomes in mouse M2

To characterize neural activity in M2 while mice engaged in the task, we simultaneously imaged the brain at cellular resolution using a two-photon microscope (Denk W et al. 1990) (**Fig. 3A**). An adeno-associated virus encoding GCaMP6s (AAV1.Syn.GCaMP6s.WPRE.SV40) was injected into layer 2/3 of M2, and a chronic glass window was implanted for optical imaging (**Fig. 3B**). GCaMP6s is a genetically encoded calcium indicator that exhibits an approximately 25% rise in fluorescence intensity per action potential in cortical pyramidal neurons (Chen TW et al. 2013). In sixteen imaging sessions, we recorded fluorescence transients from an average of 48 ± 3 neurons in layer 2/3 of M2 (mean ± *SEM*, range: 33 – 77 cells; **Fig. 3C**). All imaging was done in the right hemisphere; therefore, left and right licks were always contralateral and ipsilateral to the imaged neurons, respectively.

**Figure 3.**
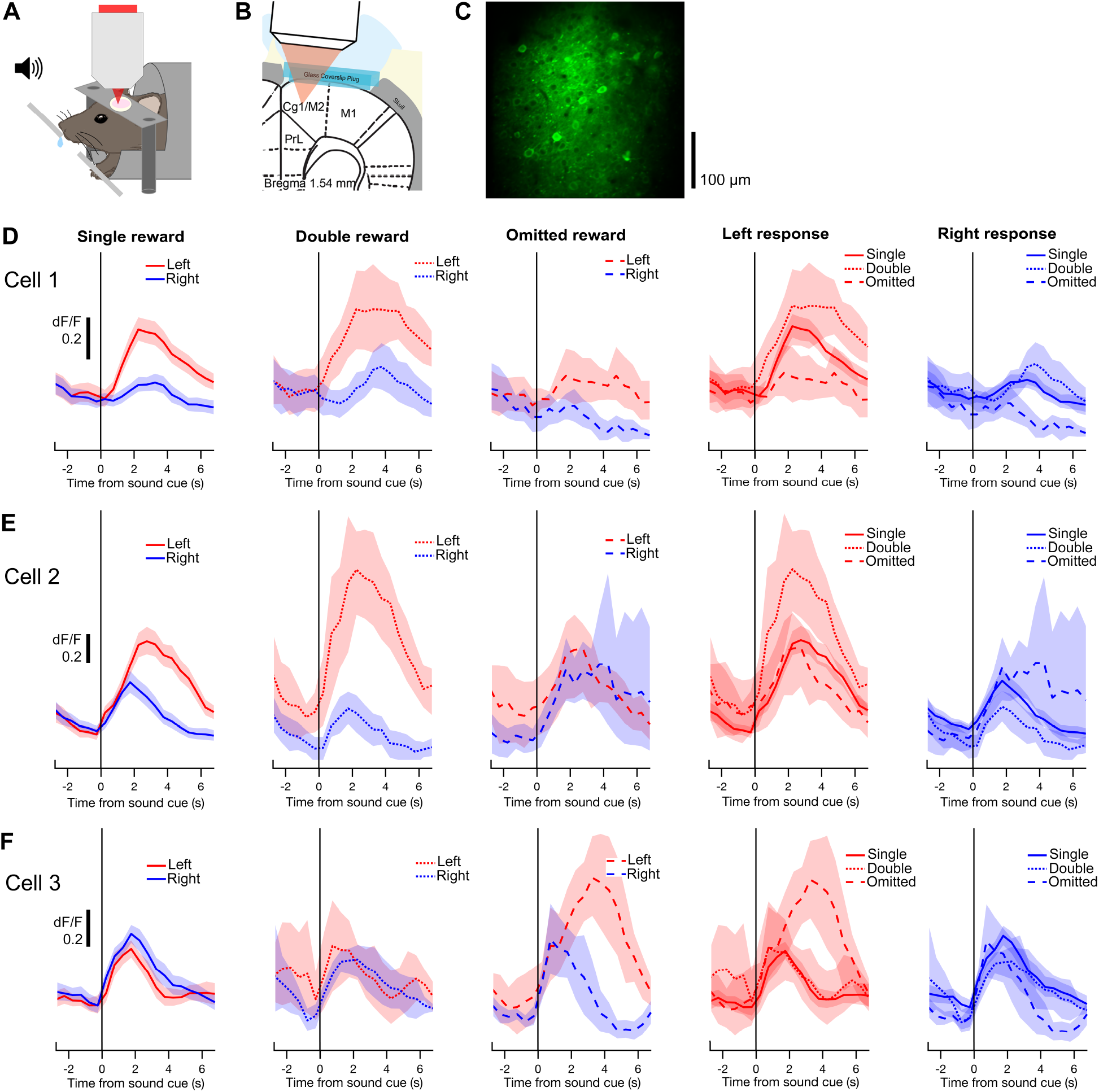
Two-photon calcium imaging of choice- and outcome-related signals in secondary motor cortex (M2). ***A***, Schematic of experimental setup for behavior with simultaneous two-photon imaging. ***B***, Schematic of preparation for *in vivo* two-photon imaging of M2. PrL, prelimbic cortex; Cg1, cingulate area 1; M1, primary motor cortex. ***C***, An example field-of-view in layer 2/3 of M2 containing GCaMP6s-expressing neurons. The image is a mean projection of the full time-lapse image stack from Experiment 13 in Table 1. ***D***, Mean fluorescence traces from an example cell, aligned to the sound cue and averaged across different subsets of trials. In the leftmost three panels, traces from left (red) and right (blue) trials are overlaid for each trial outcome. The rightmost two panels display the same data, with traces from single- (solid), double- (dotted), and omitted-reward trials (dashed) overlaid for each chosen action. Gray shading, 90% confidence interval. ***E-F***, Same as ***D*** for two additional cells.

Many M2 neurons exhibited changes in fluorescence following the sound cue, indicating task-driven neural activity. **Figure 3D** shows a neuron with preferential activity on trials in which the left spout was chosen. The activity of the same neuron was also monotonically modulated by reward size: activity was highest for double rewards, moderate for single rewards, and lowest for omitted rewards. The influence of choice and outcome on this neuron was approximately additive. However, other neurons had more complex responses. Some neurons showed a choice preference on single- and double-reward trials that was not observed on omitted-reward trials, suggesting choice-outcome interaction (**Fig. 3E**). Other neurons only exhibited significant choice preference during omitted-reward trials (**Fig. 3F**). These results highlight the diversity of choice- and outcome-related signals in M2 at the level of individual neurons.

### Persistence of choice- and outcome-related signals

To more systematically characterize choice- and outcome-related signals in M2 neurons, we used multiple linear regression analysis. For each cell, we fit a linear equation (see Materials and Methods) to estimate the dependence of its fluorescence intensity on the following predictors: choice, outcome, and the interaction of choice and outcome, for the current trial, last trial, and the trial before last (**Fig. 4A**). The analysis revealed choice-dependent activity (*p* < 0.01 for the corresponding regression coefficient in at least 5 consecutive time-bins) in a substantial fraction of M2 neurons (210/771 cells, 27%; *p* < 0.01, binomial test; **Fig. 4B**). Dependence on outcome (42 cells, 5%; **Fig. 4C**) or a choice-outcome interaction (25 cells, 3%; **Fig. 4D**) was evident in comparatively fewer cells. Notably, significant fractions of M2 neurons continued to show choice- and outcome-dependent signals well into the next trial, persisting even after the next choice was made (black bars denoting *p* < 0.01, binomial test, middle panel, **Fig. 4B-D**).

**Figure 4.**
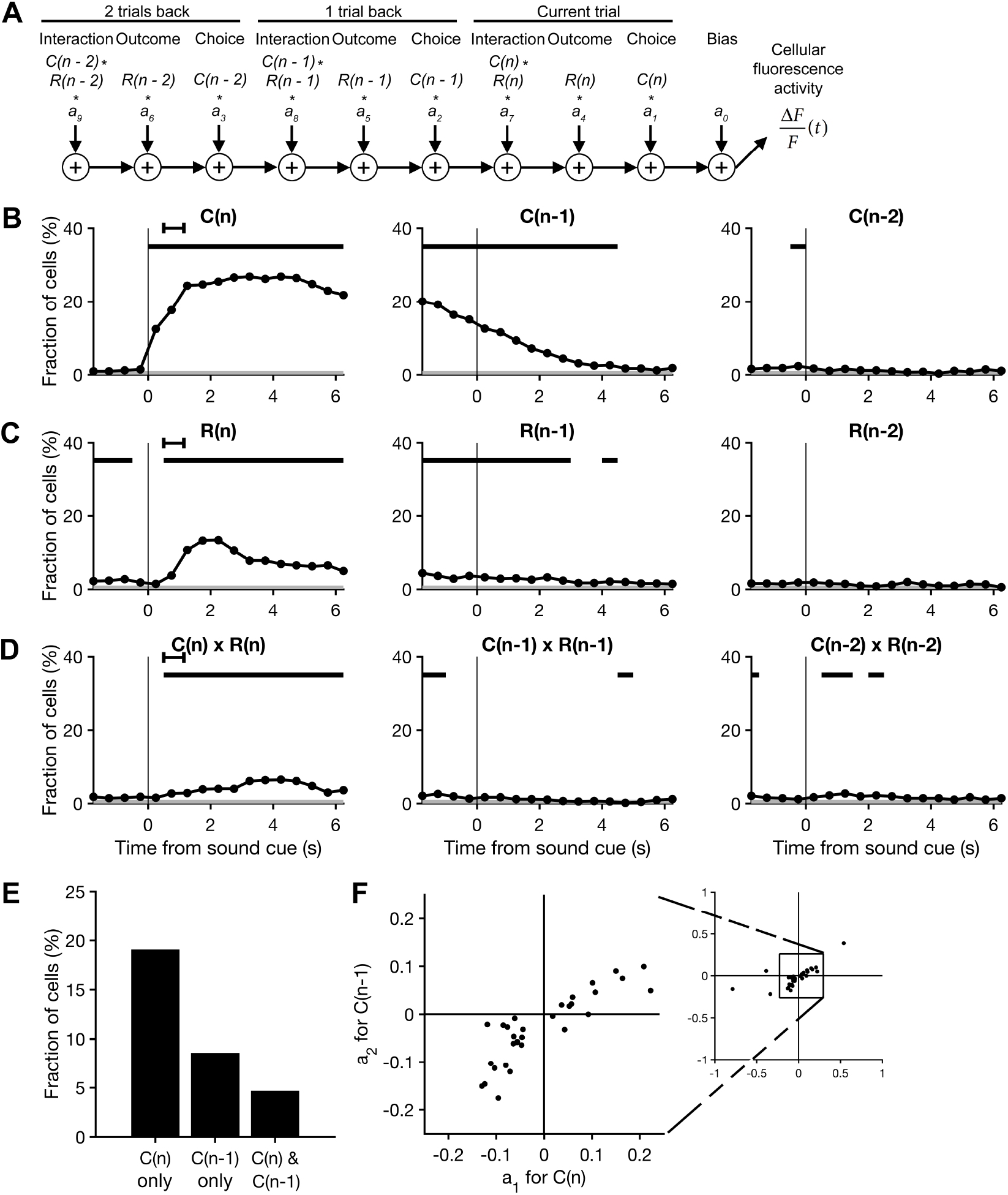
Sustained representations of choices and their outcomes in M2. ***A***, A schematic of the multiple linear regression model that was fit to the fluorescence of each neuron in each 500 ms time bin. ***B***, The proportion of cells with significant choice-dependent activity as quantified by the regression model, plotted as a function of time. The regression model accounted for the influence of choices made on the current trial (left), the last trial (middle), and the trial before last (right), as well as the additional predictors shown in ***C-D***. Significance of each predictor was tested at *α* = 0.01. Black bars, bins in which the proportion of cells with significant regression coefficients was above chance level (*p* < 0.01, binomial test). Gray shading, the significance threshold for the binomial test. Black error bar, 95% CI for time of outcome. *N* = 771 cells from 16 sessions from 10 mice. ***C-D***, Same as ***B*** for trial outcome (***C***) and the interaction of choice and outcome (***D***). ***E***, The proportion of neurons with a significant regression coefficient for the choice made in the current trial only, in the prior trial only, and in both the current and prior trials. ***F***, Scatter plot of the neurons with significant regression coefficients for both the current and prior choice. The coefficient for the current choice, *a_1_*, is plotted against the coefficient for the prior choice, *a_2_*. Right inset, the same plot expanded to show the five data points outside the range of the main axis.

The observation that M2 neurons can represent chosen actions across more than one trial led us to ask whether current and prior choices are represented at the single-unit level by the same or different populations. To address this question, we quantified the number of neurons with a significant regression coefficient only for the current choice (*p* < 0.01 for *a_1_*, the regression coefficient for the current choice estimated 2 s after cue onset, and *p* ≥ 0.01 for *a_2_*, the regression coefficient for the prior choice estimated at the time of cue onset), only for the prior choice (*p* ≥ 0.01 for *a*_1_ and *p* < 0.01 for *a_2_*), and for both the current and prior choices (*p* < 0.01 for both *a*_1_ and *a_2_*). We found that most choice-dependent M2 neurons were sensitive exclusively to the current (147/771 cells, 19%) or prior (66/771 cells, 9%) choice (**Fig. 4E**). Within the small proportion of neurons sensitive to both current and prior choices (36/771 cells, 5%), the corresponding regression coefficients were correlated and generally did not change signs (**Fig. 4F**). Therefore, the choice preferences of these cells were maintained across time-points in consecutive trials. Taken together, these results indicate that very few of the choice-selective neurons in M2 represent both the current and prior choice during the period immediately following cue onset.

### Effect of outcome on choice selectivity of individual M2 neurons

To further characterize choice representations in M2, we focused on the 226 ‘choice-selective’ neurons whose activity was found in the multiple linear regression analysis to be significantly modulated by choice, or the interaction of choice and outcome, or both. For each of these neurons, fluorescence traces were first averaged across subsets of trials according to whether the contralateral (left) or ipsilateral (right) spout was chosen (trial-averaged traces for each cell during single-reward, left trials are shown in **Fig. 5A**). A time-varying choice selectivity index was then calculated as the normalized difference between the two mean traces. During the period following cue onset in single-reward trials, the majority of choice-selective neurons preferred the contralateral choice (134/226, 59%; **Fig. 5B**). Similar to the degree of temporal variation observed across neurons in their trial-averaged activity traces, peak choice selectivity was also temporally distributed across neurons relative to the sound cue (**Fig. 5A-B**).

**Figure 5.**
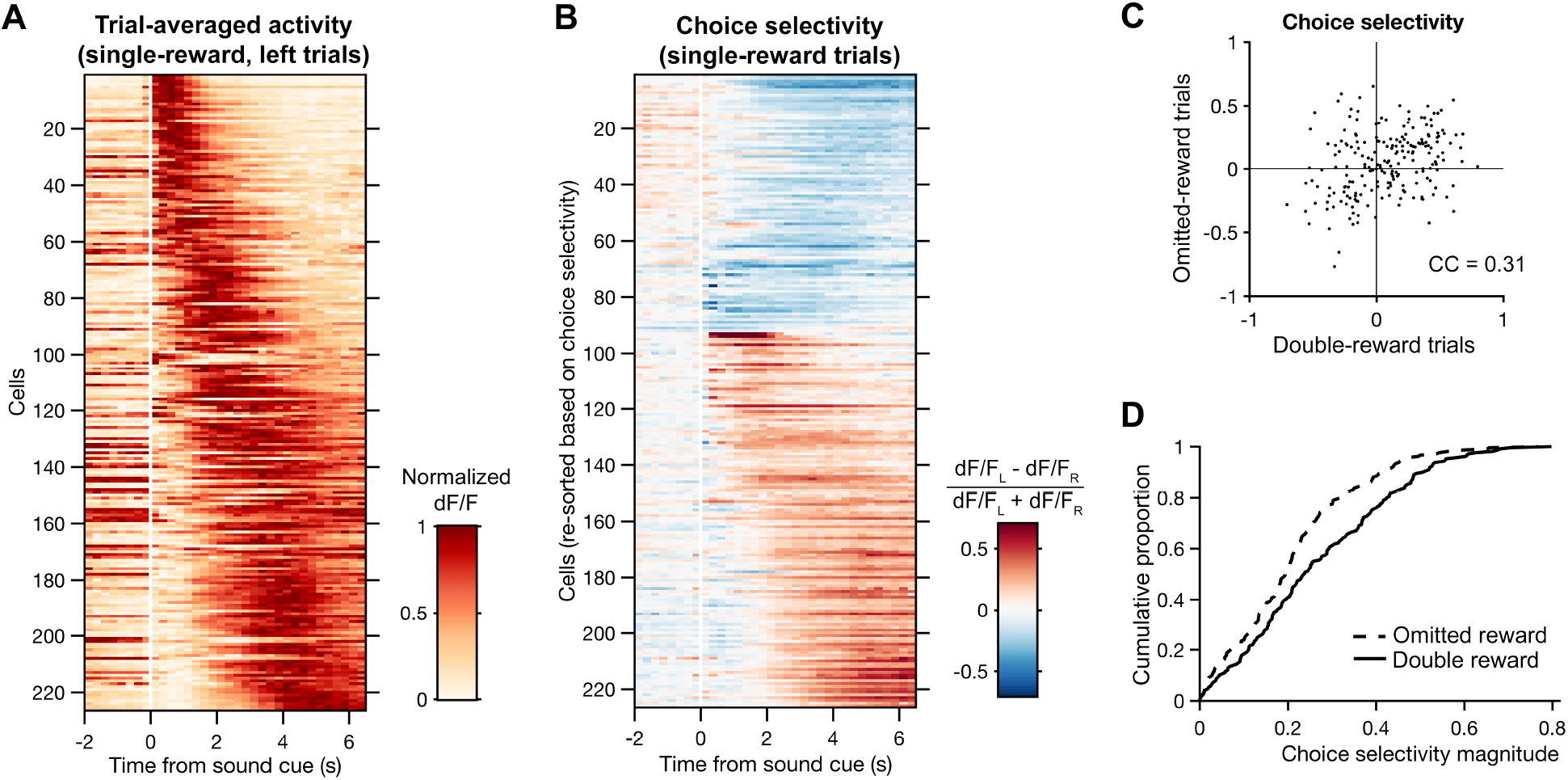
Choice representations in M2 are modified by trial outcome. ***A***, Heat map of trial-averaged fluorescence as a function of time for all choice-selective neurons during single-reward, left trials. Cells are sorted by the center-of-mass of their trial-averaged fluorescence traces. *n* = 226 cells with significant encoding of choice or an interaction of choice and outcome as determined by multiple linear regression (see Methods). ***B***, Heat map of choice selectivity for the neurons in ***A*** as a function of time during single-reward trials. Choice selectivity was calculated as the normalized difference between mean fluorescence traces from left and right trials. Red and blue shading indicate preference for left and right choices, respectively. Cells are sorted first by mean choice preference, and then by the center-of-mass of their choice selectivity traces. ***C***, Scatter plot of the neurons in ***A***, plotting the choice selectivity of each cell in omitted-reward trials against double-reward trials. CC, Pearson correlation coefficient. ***D***, Empirical cumulative distribution of choice selectivity magnitudes for double-reward (solid) and omitted-reward (dotted) trials.

Unexpected outcomes signify inaccuracies within a subject’s internal representation of the environment, including the values assigned to specific actions. Such information can be crucial to instrumental behavior. Thus, we asked whether choice representations in M2 were modified when low-probability outcomes occurred. Specifically, we examined double- and omitted-reward trials, which occurred a similar number of times per session. The choice selectivity of individual neurons clearly varied by trial type, as evidenced by the degree of scatter from the unity line in **Figure 5C**. Nevertheless, across the choice-selective neurons, the choice selectivity values for double- and omitted-reward trials were significantly correlated (correlation coefficient = 0.31, *p* = 5 × 10^−6^, *N* = 226 cells). Moreover, choice selectivity magnitudes were reduced on omitted-reward as compared to double-reward trials (*p* = 5 × 10^−5^, Wilcoxon signed-rank test; **Fig. 5D**). Extending this analysis to include all imaged neurons yielded similarly correlated choice selectivity values (correlation coefficient = 0.20, *p* = 2 × 10^−8^, *N* = 771 cells) and a similar reduction in choice selectivity magnitudes for omitted-reward relative to double-reward trials (*p* = 0.002, Wilcoxon signed-rank test). These results indicate that trial outcomes can substantially influence choice representations in M2. In particular, the absence of an expected reward in omitted-reward trials was associated with weaker representations of chosen actions at the level of single neurons.

### Reward omission weakens ensemble-level choice representations

Frontal cortical neurons often exhibit complex patterns of selectivity for task variables, and mounting evidence suggests that the associated neural representations may be best understood by focusing on the activity of large populations of neurons, rather than single units. Therefore, relative to the single unit analyses presented above, the measure of accuracy at decoding choices from ensemble activity could provide a more reliable estimate of the choice information available to downstream brain areas.

To test the effect of trial outcomes on population-level choice representations in M2, we trained linear classifiers on the ensemble activity recorded during single-reward trials, and then compared their accuracy for decoding choices across trials with different outcomes (see Materials and Methods). Baseline decoding accuracy was estimated with a Monte Carlo cross validation procedure, using ensemble activity patterns from single-reward trials as training and testing sets. For each of 30 iterations, a classifier was constructed from a random sample of 80% of these trials, and then tested on the remaining fraction. Decoding accuracy for the other outcome types was estimated by testing the same classifiers on the ensemble activity associated with the full set of double-reward, omitted-reward, and error trials.

The choices made in correct trials could be decoded from the population activity with above-chance-level accuracy at every time-point in the current trial following the animal’s response (**Fig. 6A-C, left**; black bar denotes *p* < 0.05, Wilcoxon signed-rank test, versus shuffle). An ensemble representation of the chosen action could also be detected for at least 1 s into the subsequent trial (**Fig. 6A-C, right**). For example, in single-reward trials a maximal decoding accuracy of 78 ± 2% was reached at 1.75 s after cue onset (mean ± *SEM, N* = 16 sessions; **Fig. 6A, left**). At the time of the next cue onset, accuracy remained above chance, at 63 ± 2% (mean ± *SEM, N* = 16 sessions; **Fig. 6A, right**). By contrast, in error trials the decoding accuracy increased gradually from the time of cue onset (*r* = 0.87, *p* < 0.0001, Spearman rank correlation) and only rose to significance late in the trial (**Fig. 6D, left**). It reached a maximum of 66 ± 4% at the time of the next sound cue (mean ± *SEM, N* = 16 sessions; **Fig. 6D, right**).

**Figure 6.**
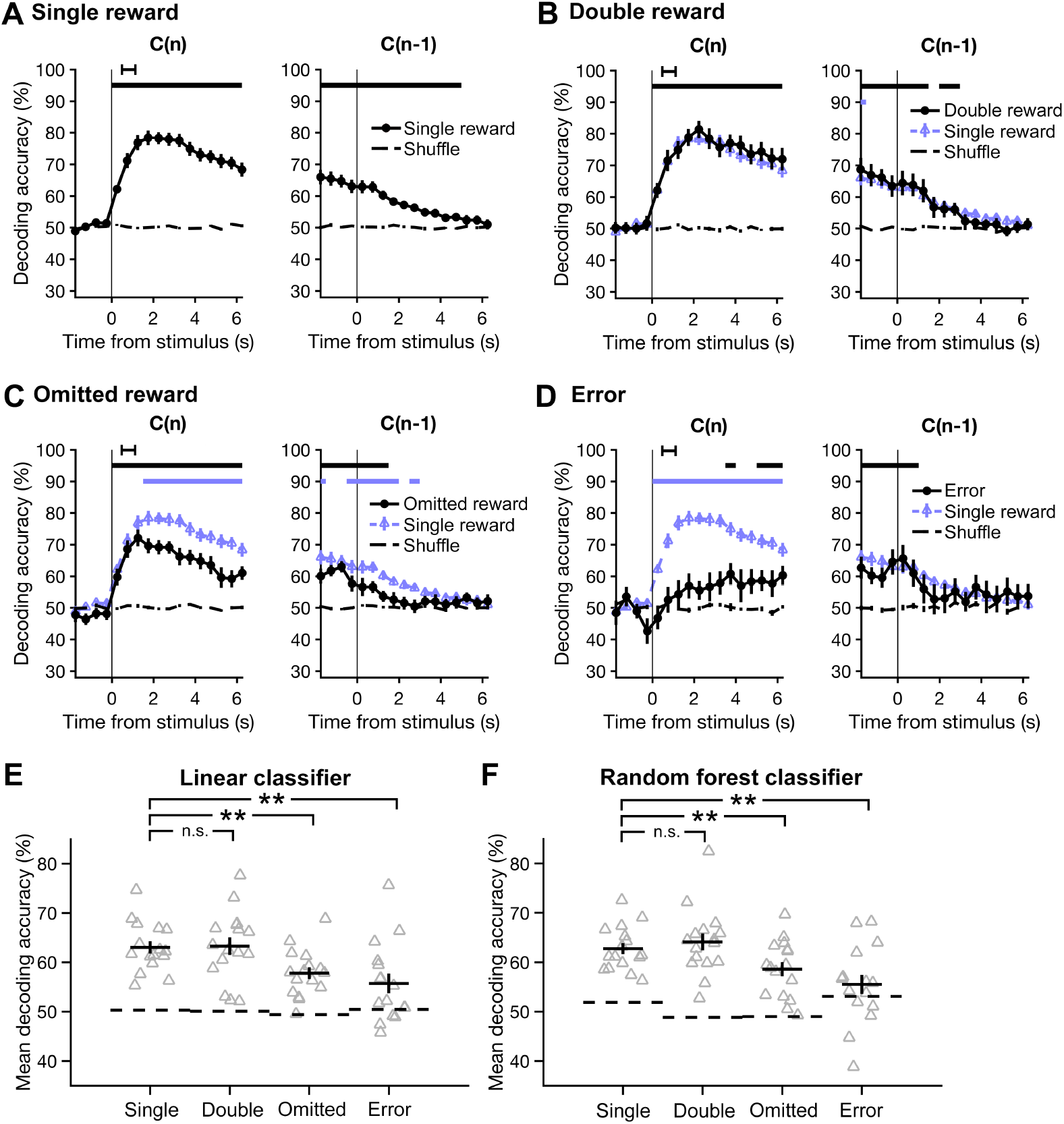
The accuracy of decoding chosen actions from the neural ensemble activity is diminished during omitted-reward and error trials. Choices were decoded using classifiers based on linear discriminant analysis, and accuracy was estimated with Monte Carlo cross-validation (repeated random subsampling). ***A***, The accuracy of decoding choices made on single-reward trials (left), or trials in which the previous outcome was single-reward (right), plotted as a function of time. Data are presented as mean ± *SEM*. Chance-level accuracy (black dashed line) was determined by testing classifiers constructed using shuffled choices. Black horizontal bars, bins significantly different from chance (*p* < 0.05, Wilcoxon signed-rank test). Black error bar, 95% confidence interval for time of outcome. ***B-D***, Same as ***A*** for double-reward (***B***), omitted-reward (***C***), and error trials (***D***). Results from single-reward trials are overlaid for visual comparison (gray triangles). Lower gray bars, bins with a significant difference in decoding accuracy relative to single-reward trials. ***E***, Mean decoding accuracy across all time-points shown in ***A-D*** for each trial outcome. Gray triangles, individual sessions. Black crosshairs, mean ± *SEM*. Wilcoxon signed-rank test: \*\**p* < 0.01; n.s., not significant. ***F***, Same as ***E*** using random forest classifiers.

For explicit comparisons of decoding accuracy across outcome conditions, we computed the mean accuracy over all time-points in the current and subsequent trial **(Fig. 6E)**. Relative to single-reward trials, mean decoding accuracy dropped significantly in omitted-reward (*p* = 0.004, Wilcoxon signed-rank test, *N* = 16 sessions) and error trials (*p* = 0.002, Wilcoxon signed-rank test, *N* = 16 sessions), with no detectable difference in double-reward trials (*p* = 0.8, Wilcoxon signed-rank test, *N* = 16 sessions).

It seems unlikely that the choice information present in M2 ensemble activity would be read out by the brain in exactly the same manner as a linear classifier. Therefore, we sought an alternative approach to determine whether the results would generalize to other methods of decoding. Specifically, we turned to random forest classification, a bootstrap aggregation method based on decision trees. Random forest classifiers operate on a fundamentally different principle than linear classifiers (see Materials and Methods), but overall, they yielded very similar results. In particular, comparisons of mean decoding accuracy again revealed marked differences between single-reward trials and omitted-reward or error trials (*p* = 0.006 and *p* = 0.003 respectively, Wilcoxon signed-rank test, *N* = 16 sessions), but no difference between single- and double-reward trials (*p* = 0.2, Wilcoxon signed-rank test, *N* = 16 sessions; **Fig. 6F**). Thus, the results of two distinct classification approaches support the same conclusion: that choice information was encoded with higher fidelity during trials in which choices were rewarded. More specifically, increases in reward magnitude (i.e., double-reward) had little impact on decoding accuracy, whereas reward absence in omitted-reward and error trials significantly diminished the accuracy with which chosen action could be decoded from neural ensemble activity patterns in M2.

### Simultaneous recording improves decoding accuracy

Our measurements of neural activity came from two-photon calcium imaging, which enabled the simultaneous acquisition of fluorescence transients from ensembles of at least 30 neurons. This allowed us to address an important, unresolved methodological question: does simultaneous recording confer a decoding advantage, relative to recording from each cell individually and then combining the data post-hoc *in silico*?

As a basis for comparison, we generated ‘pseudo-ensemble’ data for each session, in which the activity traces from each neuron were shuffled across single-reward trials in which the same action was chosen. Thus, the correspondence between individual neural activity traces and left or right choices was preserved. However, on the population level, the ensemble activity patterns no longer reflected simultaneously recorded activity. In particular, the shuffle should preserve co-fluctuations in neural activity related to the sound cue, choice, and outcome of a given trial, while disrupting the residual correlations.

To determine the extent to which correlations in neural activity were disrupted by the shuffling procedure, we first determined the correlation between each pair of neurons across trials, using the mean cellular fluorescence over the time interval from 2 to 4 s after cue onset (**Fig. 7A**). As expected, pairwise correlations were reduced within the pseudo-ensembles (**Fig. 7B**). Across all sessions, the mean Pearson correlation coefficient decreased from 0.173 ± 0.020 in simultaneously recorded ensembles to 0.022 ± 0.007 in pseudo-ensembles (*p* = 4 × 10–4, Wilcoxon signed-rank test; **Fig. 7C, left**). The mean magnitude of correlation also decreased, from 0.192 ± 0.017 in simultaneously recorded ensembles to 0.054 ± 0.005 in pseudo-ensembles (*p* = 4 × 10–4, Wilcoxon signed-rank test; **Fig. 7C, right**).

**Figure 7.**
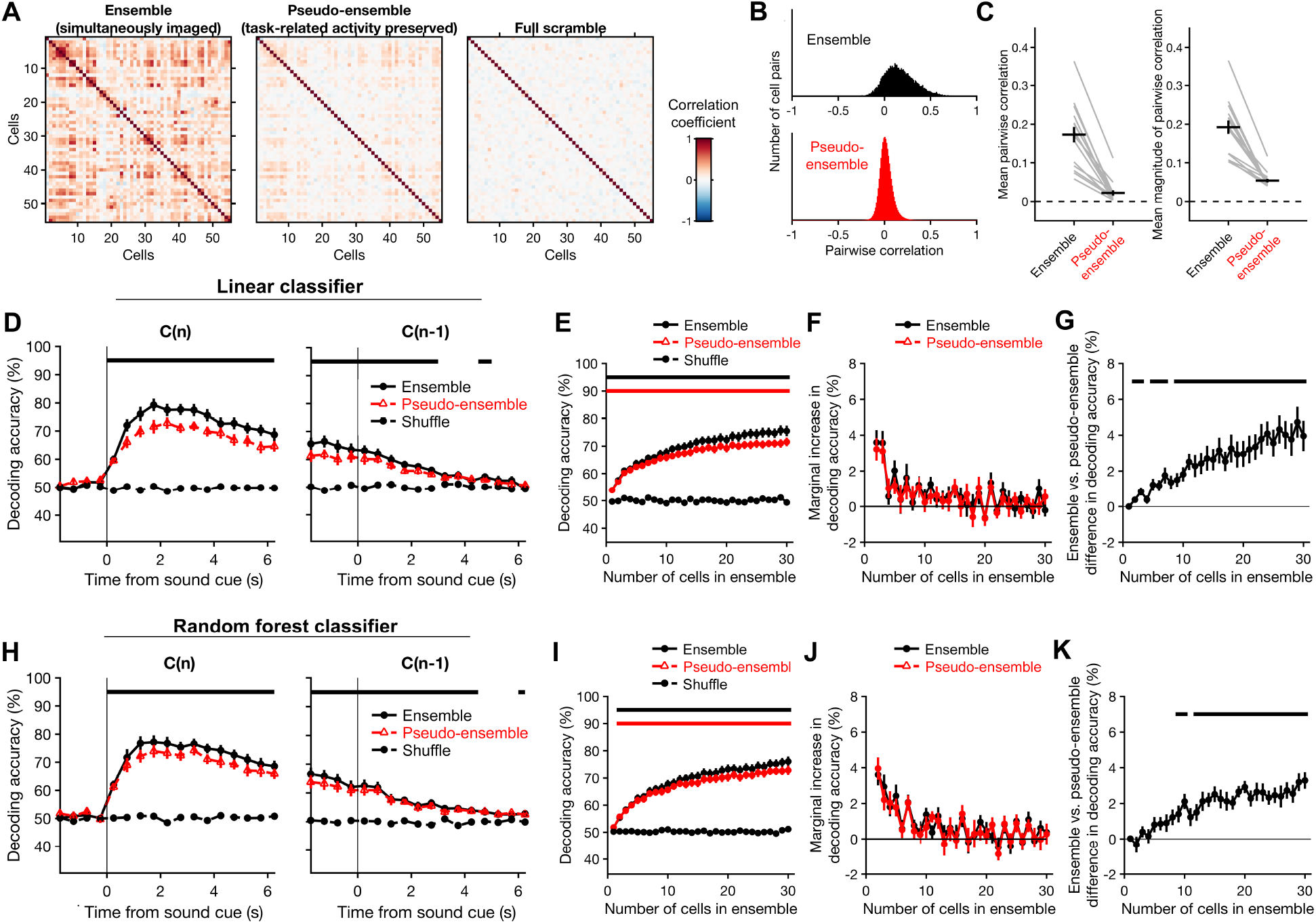
Simultaneous recording imparts a decoding advantage that increases as a function of ensemble size. The accuracy of decoding choices from the activity of simultaneously imaged ensembles of neurons was compared to that of ‘pseudo-ensembles’ in which simultaneity was disrupted by shuffling the activity traces from each neuron across trials in which the same choice was made. Only correct trials resulting in a single-reward were used for this analysis. Classification accuracy was tested using Monte Carlo cross-validation (repeated random subsampling). Chance-level accuracy was determined by testing classifiers constructed using shuffled choices. ***A***, Pearson correlation matrix for all cells from one example session, under three conditions: with simultaneity preserved (‘Ensemble’), after shuffling across trials with the same chosen action (‘Pseudo-ensemble’), and after shuffling across trials irrespective of chosen action (‘Full scramble’). Correlations were estimated from cellular fluorescence averaged over the interval from 2–4 s following cue onset. ***B***, Histogram of the Pearson correlation coefficients estimated for all pairs of cells imaged in all experiments, using simultaneous ensembles (top) and pseudo-ensembles (bottom). ***C***, Mean pairwise correlation (left), and pairwise correlation magnitude (right) across all sessions. Gray lines, means from individual sessions. Black crosshairs, grand mean ± *SEM. **D***, Performance of decoders based on linear discriminant analysis, plotted as a function of time for single-reward trials (left), or trials in which the previous outcome was single-reward (right). Accuracy of ensemble classifiers (black circles) is overlaid with that of pseudo-ensemble classifiers (red triangles) for visual comparison. Black dashed line, chance-level accuracy. ***E***, Decoder performance as a function of the number of cells used to decode the chosen action. Performance of ensemble (black circles) and pseudo-ensemble (red triangles) classifiers was estimated as the mean classification accuracy over the interval from 2–4 s following cue onset in single-reward trials. The number of cells was varied from 1–30 by drawing cells randomly from the full ensemble or pseudo-ensemble without replacement. Black dashed line, chance-level accuracy. Horizontal bars, bins in which ensemble (black, upper bar) or pseudo-ensemble (red, lower bar) classifiers performed significantly better than chance (*p* < 0.05, Wilcoxon signed-rank test). ***F***, Marginal percentage point change in decoding accuracy, plotted as a function of ensemble size for ensembles (black) and pseudo-ensembles (red). ***G***, Difference in accuracy of the ensemble and pseudo-ensemble decoders shown in ***E-F***, plotted as a function ensemble size. Black horizontal bars, bins in which the accuracy of ensemble and pseudo-ensemble classifiers differed significantly (*p* < 0.05, Wilcoxon signed-rank test). ***H-K***, Same as ***D-G*** for random forest classifiers. Data in ***D-K*** are presented as mean ± *SEM*.

**Figure 7D** shows the accuracy of decoding choices from actual and pseudo-ensemble data as a function of time during single-reward trials. To directly compare the two conditions, we again focused on the time interval from 2 to 4 s after cue onset, when decoding accuracy was highest. Over this interval, linear classifiers constructed with either simultaneous or pseudo-ensemble activity from thirty cells could decode choices with high accuracy, at 75 ± 2% and 72 ± 2%, respectively (mean ± *SEM, N* = 16 sessions; black and red bars denoting *p* < 0.01 versus shuffle, Wilcoxon signed-rank test, **Fig. 7E**). In a head-to-head comparison, decoders constructed with simultaneous activity outperformed pseudo-ensemble decoders by 4.0 ± 0.8% (mean ± *SEM, N* = 16 sessions). Random forest classifiers also exhibited a significant simultaneity effect, although it was slightly smaller than for linear classifiers. Decoding accuracy for ensemble and pseudo-ensemble activity was 76 ± 2% and 73 ± 2%, respectively, with a mean difference of 3.3 ± 0.5% in a head-to-head comparison using an ensemble size of thirty cells (mean ± *SEM*; **Fig. 7H**).

Additionally, we assessed the impact of ensemble size on decoding accuracy by training and testing classifiers using random samples of one to thirty neurons. Classifiers constructed from either ensemble or pseudo-ensemble data could decode choices with an accuracy exceeding chance for every ensemble size tested (**Fig. 7E**, black and red bars bar denoting *p* < 0.01 for ensembles and pseudo-ensembles, respectively, Wilcoxon signed-rank test). In both cases, mean decoding accuracy increased as function of ensemble size (ensembles: *r* = 0.997, *p* = 0; pseudo-ensembles: *r* = 0.996, *p* = 0; Spearman rank correlation). However, the marginal change in decoding accuracy dropped rapidly (**Fig. 7F**). For ensembles, it decreased from 3.6 ± 0.7% to 1.3 ± 0.4% after the second and tenth cell was added, respectively (mean ± *SEM; p* = 0.02, Wilcoxon signed-rank test). For pseudo-ensembles, it decreased from 3.1 ± 0.7% to 0.6 ± 0.4% (mean ± *SEM; p* = 0.02, Wilcoxon signed-rank test).

Interestingly, the decoding advantage associated with simultaneous recording was also related to ensemble size. The difference in mean decoding accuracy was significant for all ensemble sizes larger than nine cells (**Fig. 7G**, black bar denoting *p* < 0.01, Wilcoxon signed-rank test), and increased as a function of the number of cells up to the largest ensemble size examined (*r* = 0.97, *p* = 0, Spearman rank correlation). An identical analysis using random forests yielded similar results (**Fig. 7I-K**).

Taken together, our analyses reveal that choices can be decoded more accurately from simultaneously recorded population activity, relative to pseudo-ensembles in which the correlations in neural activity associated with simultaneity have been disrupted. This difference increased with the number of neurons in an ensemble, across the range of ensemble sizes tested. Moreover, the marginal decoding accuracy decreased rapidly regardless of whether ensembles or pseudo-ensembles were used—a result consistent with high levels of redundancy in the population code for chosen actions in M2.

### Discussion

How does the outcome of a chosen action influence how it is represented in the brain? In this study, we used a two-choice discrimination task with probabilistic outcomes to investigate this question in the M2 region of the murine medial frontal cortex (MFC). The results help to illuminate how information related to choices and their outcomes are integrated within the frontal lobe. M2 neurons were found to robustly encode rewarded choices; however, choice-related signals diminished when a rewarding outcome was omitted. Furthermore, an increase in the magnitude of reinforcement had far less impact on choice representations than did its categorical presence or absence. The preferential encoding of rewarded choices in M2 provides a plausible mechanism that may underlie its established role in the learning and implementation of reinforced actions during instrumental behavior.

### Cortical representation of prior choices in rodents

Optimal performance in the discrimination task requires the subject to choose strictly based on auditory cues. In principle, information about past actions could be discarded or ignored. Therefore, in these well-trained animals it was somewhat surprising to observe robust and persistent choice representations, both at the level of single neurons (**Figs. 4–5**) and ensembles (**Figs. 6–7**). However, similar task-irrelevant information coding has been reported elsewhere—in the monkey prefrontal cortex (Genovesio A et al. 2014), as well as in the posterior parietal cortex of rodents (Morcos AS and CD Harvey 2016). At the behavioral level, response biases based on choice and outcome history have been observed in human subjects during perceptual tasks even after extensive training (Frund I et al. 2014; Abrahamyan A et al. 2016). Our findings and previous results therefore suggest that under some circumstances, higher-order cortical areas continue to monitor past choices and outcomes, even if task performance does not strictly require such information.

Previous studies have reported that neurons in the rodent MFC encode past choices (Sul JH *et al*. 2011; Siniscalchi MJ *et al*. 2016) and their outcomes (Kargo WJ et al. 2007; Sul JH *et al*. 2011; Yuan Y et al. 2015). However, sustained choice-related signals are not unique to this brain region. They have also been found in other nodes of the frontal-striatal network including the dorsomedial striatum (Kim H et al. 2013) and orbitofrontal cortex (Sul JH *et al*. 2010), as well as the posterior parietal cortex (Hwang EJ et al. 2017). This is not to say that all cortical regions exhibit persistent signals associated with chosen action. For example, our previous study detected only very brief choice signals in the mouse primary visual cortex (Siniscalchi MJ *et al*. 2016). Similarly, choice signals like those found in dorsomedial striatum lasted only transiently in dorsolateral striatum (Kim H *et al*. 2013). This point is notable because M2 and other medial frontal areas send dense projections to dorsomedial striatum, while afferents in dorsolateral striatum come mostly from primary motor cortex (Reep RL and JV Corwin 1999).

The extended time course of the choice signals we observed could serve as an eligibility trace that keeps recently performed actions available for learning. In the current study, the representation of choice-related information in M2 neurons persisted into the middle of the next trial (**Fig. 6A**). It is worth noting that our previous study detected significant choice-related signals over an even longer duration—up to two trials after the corresponding action was chosen (Siniscalchi MJ *et al*. 2016). Interestingly, the prior study employed a rule-switching task in which subjects were required to monitor choices and their outcomes. This raises an intriguing possibility: that the temporal scale of choice history signals may depend on the task demands (Bernacchia A et al. 2011; Donahue CH *et al*. 2013). Namely, the optimal learning rate depends on the volatility of the environment (Behrens TE et al. 2007; Farashahi S et al. 2017). If the persistence of choice representations is indeed a flexible parameter, then it could allow the system to adapt to changes in volatility by serving as a point of adjustment for the temporal integration of choice information.

### Enhanced population coding for rewarded choices

Our results reveal that neural representations of chosen actions in mouse M2 are outcome-dependent. This finding agrees in principle with a previous study demonstrating that rewarded choices are more reliably encoded relative to unrewarded choices in the primate supplementary eye field and dorsolateral prefrontal cortex during a matching-pennies task (Donahue CH *et al*. 2013). Interestingly, the effect was not evident in recordings from the same set of neurons during a visual search task in which a visuospatial cue instructed the correct response at the beginning of each trial. The critical difference between the two tasks may be the presence of an instructive cue—a feature which the visual search task shares with the task used in our current study. Notably, we did find a robust reward-associated enhancement of choice coding under these circumstances. One possible explanation is that, in contrast to visuospatial instruction, the cues used in our auditory discrimination task only came to be associated with the correct choice through learning. Therefore, the reward-associated enhancement we observed may reflect an action-monitoring process associated with the maintenance of arbitrary sensorimotor associations—a process which may be unnecessary during the less demanding visual search task. Another possible explanation regards our use of ensemble decoding, which may be more sensitive than decoding from single units and can thus be expected to detect smaller differences between outcome conditions.

In general, the neural correlates of choice and outcome history have been studied using binary outcome conditions in which a reward is either provided or withheld on a given trial. We sought to extend the results of these prior studies by comparing effects of multiple reward magnitudes. Additionally, we were able to dissociate effects of performance and outcome by measuring the impacts of unexpected reward omissions and windfalls in mice trained to a very high level of proficiency (>90% accuracy).

At the behavioral level, reward size affected consummatory licking as well as the likelihood of a response to the next cue, but failed to influence the accuracy of responses **(Fig. 2A-C)**. This suggests that infrequent changes in reward magnitude impacted motivation without significantly influencing choices. Moreover, the neural ensemble representations of chosen actions did not differ between single- and double-reward trials. Instead, the greatest contrast was found between rewarded and unrewarded choices. Our results therefore suggest that the influence of outcome on choice representations in M2 is driven less by the magnitude of reward than by its explicit presence or absence. However, because actions were reinforced immediately in our task design, it remains possible that floor or ceiling effects could have limited the outcome sensitivity of the choice-selective population.

What physiological mechanisms underlie the outcome dependence of choice signals observed in M2? One intriguing possibility concerns the role of neuromodulation, which may directly reconfigure the local network dynamics, or act on inputs to M2. In particular, dopaminergic (Schultz W et al. 1997) and cholinergic (Hangya B et al. 2015) neurons are known to carry signals related to reward. Furthermore, reward-dependent activation of dopaminergic projections to nearby primary motor cortex have been implicated in motor skill learning (Hosp JA et al. 2011; Leemburg S et al. 2018). It is therefore interesting to speculate on whether similar mechanisms might contribute to associative learning (Takehara-Nishiuchi K and BL McNaughton 2008) and more specifically, to the auditory-motor associations necessary for performance of the task presented here. In any case, the impact of neuromodulators on motor cortical choice signaling will comprise an exciting topic for future research.

### Persistent neural and behavioral effects associated with errors

The actions chosen on error trials were decoded least accurately from the corresponding ensemble activity **(Fig. 6E-F)**. This result is consistent with an earlier study that revealed disrupted MFC ensemble representations for choices and their outcomes during periods when rats committed multiple errors in a radial arm maze (Lapish CC et al. 2008; Hyman JM et al. 2013). The discretized trial structure of our auditory discrimination task allowed us to build upon this prior result by measuring the reliability of ensemble representations into the next trial. Furthermore, the inclusion of omitted-reward trials allowed direct comparisons of ensemble representations associated with correct and incorrect choices, independent of the associated outcomes.

Another previous study demonstrated a tight relationship between sustained error signals in MFC and behavioral performance in the next trial, measured as post-error slowing during a timing task (Narayanan NS et al. 2013). Similarly, our analysis revealed not only an error-related decrement in the fidelity of neural choice representations, but also a behavioral performance decrement following error trials that could not be explained by reward omission alone. These results may provide some insight into the sources of error for these well-trained subjects. Specifically, the prolonged time course of the neural and behavioral effects associated with errors suggests that they may have arisen in part due to factors that spanned multiple trials—such as periods of hypo- or hyper-arousal. In any case, these results together with prior studies indicate that errors are often associated with persistent internal states that can impact subsequent behavioral performance.

### Simultaneous recording confers a modest decoding advantage in M2

Prior theoretical work has demonstrated that correlated variability in neural populations can either degrade or enhance population coding, depending on the interaction between signal and noise correlations (Averbeck BB and D Lee 2003; Averbeck BB et al. 2006). The analysis shown in **Fig. 7** revealed that chosen actions could be decoded more accurately from simultaneously recorded ensembles, relative to pseudo-ensembles in which the correlations in neural activity associated with simultaneity (i.e., noise correlations) had been disrupted. In particular, the observed effect of simultaneity seems to have resulted from the preservation of largely positive correlations in trial-to-trial neural variability unrelated to the chosen action (**Figs. 7B-C**). Possible sources of noise correlations in our recordings could include unobserved behavior such as whisking, features of the network architecture, or changes in internal state associated with motivation or arousal.

The effect of simultaneity increased with the number of neurons in an ensemble, across the range of ensemble sizes tested (**Fig. 7G**). Notably, an earlier study in the primate supplementary motor area found no statistically significant effect of correlated spike-count variability on the encoding of movements by ensembles of 3–8 neurons (Averbeck BB and D Lee 2006). Our analyses only revealed a consistent simultaneity effect for ensembles larger than nine neurons, which may highlight the utility of large-scale recordings for addressing this question. However, it should be emphasized that even for ensembles of 30 cells, the comparative advantage for ensembles was modest (4%), and choices could still be decoded from pseudo-ensembles with above-chance-level accuracy at every ensemble size. Furthermore, estimated correlations between neurons tend to strengthen at longer time scales (Averbeck BB and D Lee 2003). Hence, the wider time-bins used in our study (500- vs. 66 ms), as well as the slower dynamics associated with calcium imaging could explain why our analyses were more sensitive to correlated variability. We also found that marginal decoding accuracy decreased rapidly as cells were added to the population, for both ensembles and pseudo-ensembles (**Figs. 7F, 7J**). This result suggests a high level of redundancy in M2 population codes, similar to previous results found in the rat primary motor cortex during a simple reaction time task (Narayanan NS et al. 2005).

### Insights into the role of M2 in goal-directed behavior

The choice selectivity magnitudes of individual neurons (**Fig. 5D**) and the accuracy of choice decoding from ensemble activity (**Fig. 6E-F**) both decreased from double- to omitted-rewarded trials, and then further decreased in error trials. How do the observed physiological changes ultimately impact behavior? Causal perturbations aimed at addressing this question will require a more detailed understanding of how genetically (Kvitsiani D et al. 2013; Pinto L and Y Dan 2015; Kamigaki T and Y Dan 2017) or anatomically identified subtypes of frontal cortical neurons (Li N et al. 2015; Chen TW et al. 2017; Otis JM et al. 2017) contribute to the choice signals observed in our experiments.

Goal-directed behavior requires the capacity to adjust the current policy for action selection according to the impact of past choices on the likelihood of a desired outcome. Our results demonstrate that sustained neural representations of chosen actions in mouse M2 are sensitive to their resultant outcomes, such that rewarded choices are more robustly encoded. In turn, the preferential encoding of rewarded choices could allow the frontal cortex to bias the influence of recent, positively reinforced actions on future decisions. This proposed mechanism would help to explain effects of lesioning (Passingham RE et al. 1988; Gremel CM and RM Costa 2013) and inactivation (Siniscalchi MJ *et al*. 2016; Makino H *et al*. 2017) that have implicated M2 more broadly in the learning and implementation of voluntary behavior. In summary, our results contribute to a growing body of evidence supporting a role for MFC, and M2 more specifically, in the flexible execution of goal-directed actions.

## Acknowledgements

This work was supported by National Institute of Mental Health grant R01MH112750 (A.C.K.) and R21MH118596 (A.C.K.), National Institute on Aging grant P50AG047270 (A.C.K.), SFARI Explorer Award (A.C.K.), National Institutes of Health training grant T32NS041228 (M.J.S.), National Science Foundation Graduate Research Fellowship DGE-1122492 (M.J.S.), and China Scholarship Council-Yale World Scholars Fellowship (H.W.).

## Author contributions

M.J.S. and A.C.K. designed the experiments. M.J.S. conducted the experiments. M.J.S., H.W., and A.C.K. analyzed the data. M.J.S. and A.C.K. wrote the paper.

